# A novel human fetal lung-derived alveolar organoid model reveals mechanisms of surfactant protein C maturation relevant to interstitial lung disease

**DOI:** 10.1101/2023.08.30.555522

**Authors:** Kyungtae Lim, Eimear N. Rutherford, Dawei Sun, Dick J. H. Van den Boomen, James R. Edgar, Jae Hak Bang, Lydia E. Matesic, Joo-Hyeon Lee, Paul J. Lehner, Stefan J. Marciniak, Emma L. Rawlins, Jennifer A. Dickens

**Author notes:** co-first authors (Kyungtae Lim, Eimear N. Rutherford).

## Abstract

Alveolar type 2 (AT2) cells maintain lung health by acting as stem cells and producing pulmonary surfactant^1–3^. AT2 dysfunction underlies many lung diseases including interstitial lung disease (ILD), in which some inherited forms result from mislocalisation of surfactant protein C (SFTPC) variants^4,5^. Disease modelling and dissection of mechanisms remains challenging due to complexities in deriving and maintaining AT2 cells *ex vivo.* Here, we describe the development of expandable adult AT2-like organoids derived from human fetal lung which are phenotypically stable, can differentiate into AT1-like cells and are genetically manipulable. We use these organoids to test key effectors of SFTPC maturation identified in a forward genetic screen including the E3 ligase ITCH, demonstrating that their depletion phenocopies the pathological SFTPC redistribution seen for the SFTPC-I73T variant. In summary, we demonstrate the development of a novel alveolar organoid model and use it to identify effectors of SFTPC maturation necessary for AT2 health.

---

Despite its role in increasing the stability of pulmonary surfactant, SFTPC is not absolutely required for lung development and surfactant secretion^5^. However, its aberrant handling during intracellular trafficking and maturation results in toxic gain-of-function effects. This is demonstrated by the pathogenic variant I73T, which accumulates immature isoforms at the plasma membrane and causes heritable forms of pulmonary fibrosis^6,7^. Study of this variant in immortalised cells has suggested SFTPC trafficking into multivesicular bodies (MVBs) is indirect (via the plasma membrane) and requires its ubiquitination^8^. Despite the substituted amino acid (I73) lying on the opposite side of the membrane to the ubiquitinated site at K6, failure of ubiquitination appears to be the key determinant of SFTPC-I73T redistribution and the cause of AT2 dysfunction and consequently ILD. Immortalised cells can be used to generate hypotheses, but questions regarding mechanisms and key effectors of SFTPC trafficking can ultimately only be answered using genetically-manipulable physiological AT2 cells which endogenously process surfactant.

AT2 organoids have been grown from human adult lungs and used as models of SARS-CoV-2 infection^9–11^. Human adult AT2 organoids proliferate slowly and are difficult to genetically manipulate. AT2 organoids can also be derived from pluripotent stem cells (PSC-iAT2s)^12–14^. These cells can readily be genetically manipulated, but their differentiation is complicated, taking more than 30 days^15^, and PSC-iAT2s can spontaneously dedifferentiate to other organ lineages^14^. AT2 cells have been derived from human fetal lungs, but these cells could neither be genetically manipulated nor maintained long-term in culture^16^. A complementary method for growing genetically-manipulable human AT2 cells would therefore be of great value for investigating surfactant trafficking and lung disease.

During human lung development the distal tip epithelial cells act as multipotent progenitors^17,18^. From ∼15 pcw (post conception weeks) the human tips retain progenitor marker expression, but also upregulate markers of AT2 cells^19^. Immature AT2 cells appear in the tissue from 17 pcw^19,20^. We have recently shown that 16-22 pcw epithelial tip cells can be expanded as organoids and differentiated to AT2 cells^19^. However, the differentiated AT2 organoids were not proliferative, limiting their use for functional studies and genetic manipulation. We have now developed a highly robust, efficient, and scalable culture condition that induces the differentiation of 16-22 pcw fetal lung tip cells into mature AT2 cells which grow as expandable 3D organoids. Here, we characterise the fetal-derived AT2 organoids (hereafter fdAT2), showing that they are stable over long-term passaging, efficiently process and secrete surfactant, and can differentiate into AT1-like cells *in vitro* and in mouse lung transplantation assays. We use a forward genetic screen to identify candidate effectors of SFTPC trafficking which we validate using CRISPR interference (CRISPRi) in the fdAT2 organoids. We demonstrate that trafficking of SFTPC requires ubiquitination by HECT domain E3 ligases, particularly ITCH, and that their depletion phenocopies the redistribution seen for the pathological SFTPC^I73T^ variant.

Our AT2 medium directly induces the differentiation of 16-22 pcw fetal lung tip cells into mature AT2 cells which grow as expandable 3D organoids (Fig. 1A; Extended Data Fig. 1A-C). The AT2 organoids form within 3 weeks and can be split every week, for over 20 passages, while sustaining *SFTPC* promoter-GFP reporter and AT2 marker expression (Fig. 1B; Extended Data Fig. 1D,E). They retain these characteristics following cryopreservation and thawing. The fetal-derived AT2 (fdAT2) organoids show mature AT2 cell features including, genesis of lamellar bodies and production and secretion of mature forms of SFTPB and SFTPC (Fig. 1C-D; Extended Data Fig. 1F). The organoids express proteins required for surfactant production, such as LAMP3, ABCA3, and NAPSA, as well as typical AT2 markers HTII-280 and HOPX, whereas a fetal tip progenitor marker, SOX9 was not detected (Fig. 1E). Immature SFTPC is detectable at the plasma membrane in these cells, supporting our previous model of SFTPC trafficking (Fig. 3C) in which proprotein transits the cell surface before it is endocytosed and cleaved *en route* to later compartments^8^ (Fig. 1F). In addition, the fdAT2 organoids expressed cellular polarity markers ZO-1 and laminin, and proliferation marker, Ki67 (Fig. 1E). Proliferation of the fdAT2 organoids was greatly reduced when FGF7 was removed from the medium (Extended Data Fig. 1G-I), suggesting that FGF7 is a key mitogen for these cells, consistent with recent mouse data^21^. These culture conditions induce the differentiation of 16-22 pcw lung epithelial progenitors into mature AT2 cells which maintain identity and function during prolonged passaging.

**Fig. 1.**
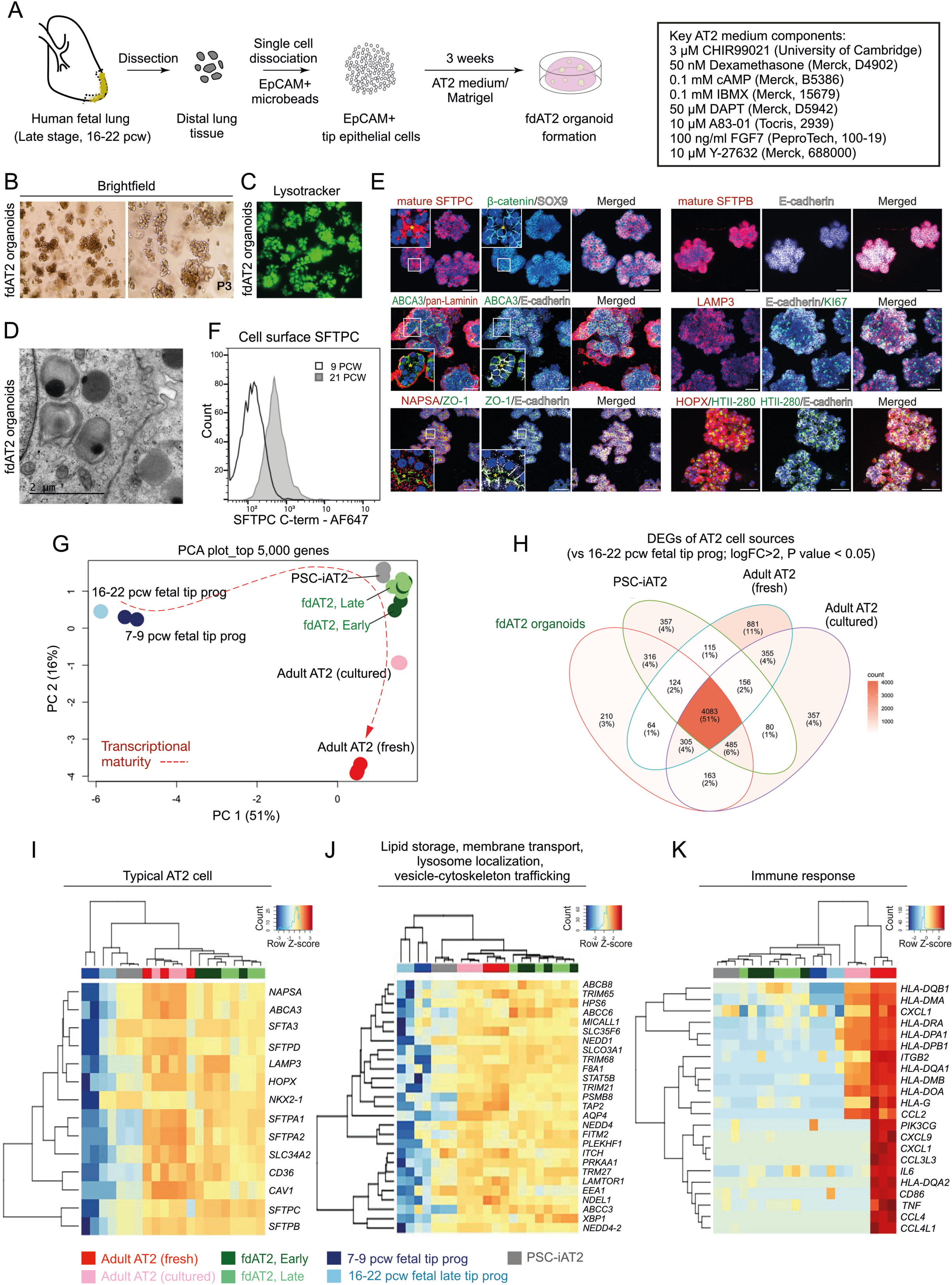
Human fetal-derived alveolar type 2 organoids show similar surfactant protein production, trafficking, and secretion to adult AT2 cells. (A) Experimental scheme and culture medium for derivation and establishment of fdAT2 organoids from human fetal lungs at 16-22 pcw. (B) Bright-field images of two independent fdAT2 organoid lines at P3, which were established from 19 pcw fetal lung tissue. (C) Uptake of lysotracker green DND-26 that stains acidic compartments within an fdAT2 organoid line at P14, showing the accumulation of lamellar bodies (acidic lysosome-related organelles). (D) Electron microscopy showing the presence of lamellar bodies with characteristic concentric lamellar membranes. Scale bar, 2 μm. (E) Immunofluorescence images of fdAT2 organoids using surfactant protein production-associated markers including mature SFTPC and SFTPB, LAMP3, NAPSA, and ABCA3, plus typical alveolar type 2 cell-lineage markers HTII-280 and HOPX, epithelial cell polarity markers E-cadherin, pan-Laminin, and ZO1, and a proliferation marker, KI67. *Asterisk (yellow) indicates apical lumen. Organoids were at p10-15. DAPI (blue), nuclei. Scale bars, 50 μm. (F) Flow cytometric analysis of cell surface proSFTPC in 9 pcw lung tip progenitor organoids and 21 pcw fdAT2 organoids as measured by C-terminal SFTPC antibody which recognises the full-length protein. (G) Principal component analysis (PCA) plot of transcriptomic profiles of the fdAT2 organoids at early and late passages (fdAT2; Early and Late), lung tip progenitor organoids from 7-9 pcw and 16-22 pcw, and other alveolar type 2 cells that were previously reported, PSC-iAT2, adult AT2 cells cultured or freshly isolated from adult human lung^2,3^. (H) Venn diagram illustrating the number and the proportion of unique or shared genes from AT2 cells of different sources. The genes for each AT2 cell type that were differentially expressed compared to fetal 16-22 pcw tip progenitor organoids were included (log_2_FC > 2, *P*-value < 0.05; Extended Data Table 2). (I-K) Heatmap of DEGs associated with typical AT2 cell fate (I), trafficking (J), and immune response (K). A list of trafficking-related genes was selectively obtained following gene ontology (GO) analysis for the GO terms; lipid storage, membrane transport, lysosome localisation, vesicle-cytoskeleton trafficking; see also Extended Data Fig. 2C.

To further investigate the maturity of fdAT2 organoids, we compared the transcriptome of four independent lines at early passage (fdAT2 early; at P1,4,6,7) and late passage (fdAT2 late; at P12,13,16,17) with those of pluripotent stem cell-derived 3D cultured induced AT2 (PSC-iAT2^12^), freshly-isolated and cultured adult AT2 cells^13^ and 8-9 pcw and 16-20 pcw fetal lung tip progenitor organoids^19^ (Fig. 1G). Our fdAT2 organoids clustered with other cultured AT2 cells, but were distinct from the fetal lung tip progenitors (PC1, Fig. 1G; Extended Data Fig. 2A). The fdAT2 organoids were transcriptionally closest to the PSC-iAT2 (Fig. 1G). The cultured adult AT2 cells sat between the fdAT2 organoids and the freshly-isolated adult AT2 cells, indicating a culture-effect on the gene expression profile of adult AT2 cells^22^ (Fig. 1G). A direct comparison of the fdAT2s and adult cultured AT2s revealed that the most significantly increased transcripts in fdAT2s are related to cell division (Extended Data Fig. 2B), consistent with their expandability. Importantly, the transcriptional profile of the fdAT2 organoids remained stable during the extended culture period, suggesting that expansion of these organoids does not affect AT2 cell identity/function (Fig. 1G; Extended Data Fig. 2A,B).

To further characterise the fdAT2 organoids, we extracted AT2-specific differentially expressed genes (DEGs) by comparing AT2 cells from each source to 16-20 pcw fetal tip progenitor organoids (Fig. 1H; log_2_FC>2, *P*-value<0.05). Comparison of these DEGs identified gene expression profiles that distinguish the different AT2 cells. This showed that 51% of DEGs (4,083) are commonly shared across all AT2 cell sources (Fig. 1H; centre). Gene ontology (GO) analysis showed that these genes are related to cytoplasmic translation, protein transport, vesicle-mediated transport, and ER-Golgi vesicle-mediated transport (Extended Data Fig. 2C). Consistent with this, the fdAT2 organoids exhibited a gene expression profile related to surfactant protein synthesis (Fig. 1I). Our DEG analysis also identified subsets of genes that were partially shared, or uniquely expressed, in the different AT2 cells (Fig. 1H; Extended Data Fig. 2C). For example, GO analysis for 532 DEGs that are shared by fdAT2 organoids and adult AT2 cells, but not by PSC-iAT2, showed terms associated with vesicle cytoskeletal trafficking, lipid storage, transmembrane transport, and lysosome localization (Fig. 1H,J; Extended Data Fig. 2C’). These GO terms are strongly correlated with the physiological surfactant-producing function of the AT2 cells (Fig. 1D-F) highlighting the utility of the fdAT2 organoids to study surfactant processing. By contrast, the fetal-derived and iPSC-derived AT2 organoids were missing 355 DEGs related to antigen processing and presentation via MHC class II and immune response that only the adult AT2 cells expressed (Fig. 1H,K; Extended Data Fig. 2C’’), suggesting that immune function cannot be acquired in a cell-autonomous manner *in vitro*.

Next, we investigated how far the transcriptional state of the fdAT2 organoids resembles the PSC-iAT2. Their transcriptome is broadly similar (Fig. 1G, Extended data Fig. 2A). However, a direct comparison indicated >2000 DEGs (Extended Data Fig. 2D). Gene set enrichment analysis (GSEA) showed that the fdAT2 organoids were enriched with gene sets associated with surfactant metabolism compared to PSC-iAT2, although they share many pathways (Extended Data Fig. 2E-G). Overall, these data confirm that fdAT2 organoids strongly resemble adult AT2 cells at a transcriptional level and possess the capacity for mature surfactant protein production, trafficking, and secretion, while proliferating. However, the fdAT2 organoids lack immune response-related features. This may reflect some immaturity, or, more likely, the sterile environment in which they are grown; supported by the observation that adult AT2 gradually lose their immune signature when cultured (Fig. 1K, Extended Data Fig. 2A).

**Fig. 2.**
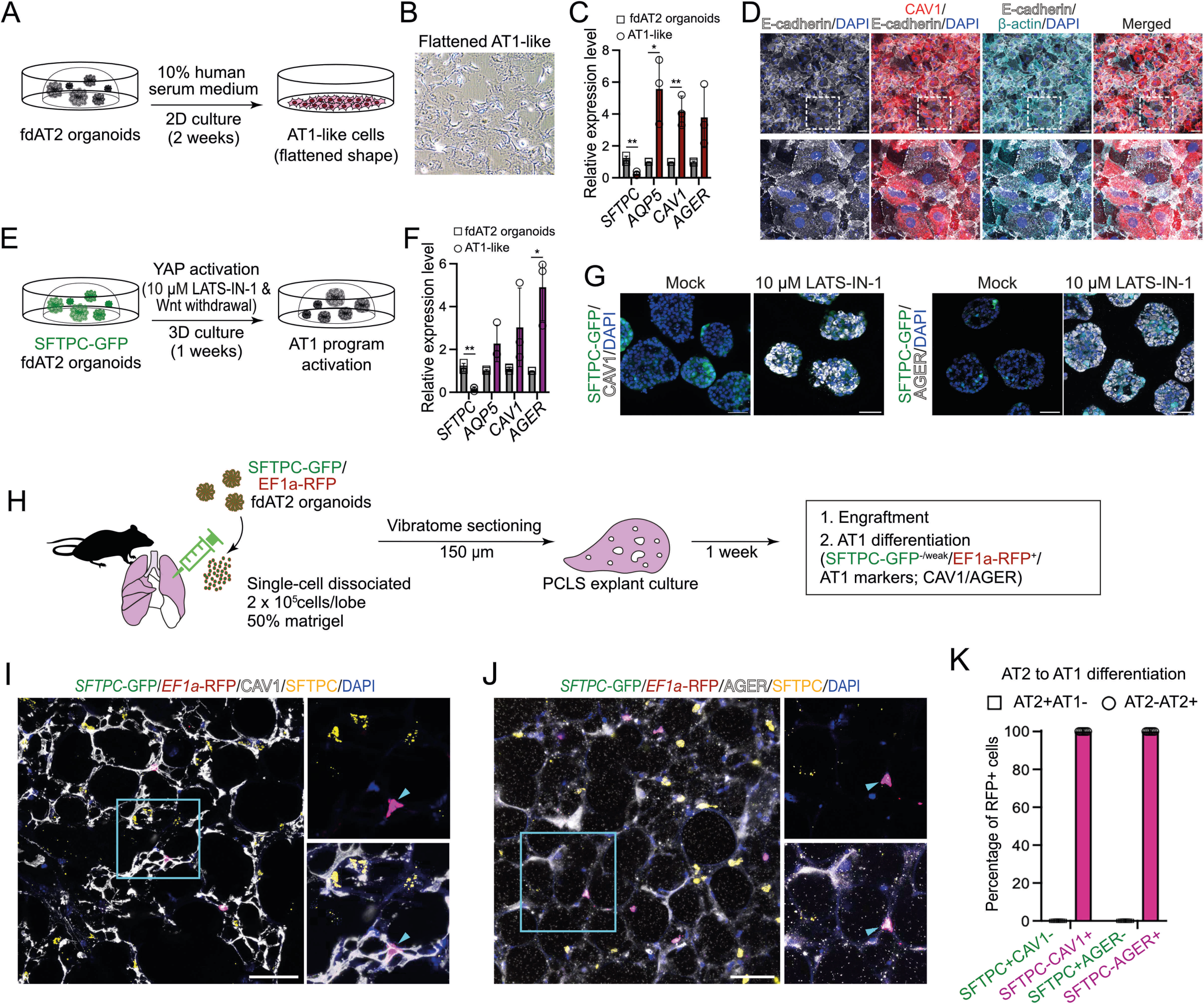
Alveolar type 1 cell fate differentiation of AT2 organoids under *in vitro* and *ex vivo* culture conditions. (A-D) Alveolar type 1 cell (AT1) fate differentiation of fdAT2 organoids in 10% human serum-containing medium on 2D culture. Experimental scheme of AT1 differentiation (A), a bright-field image of AT2 organoids upon AT1 differentiation (B), qRT-PCR analysis of AT1 markers *AQP5*, *CAV1*, and *AGER*, and an AT2 marker, *SFTPC* (C), and immunofluorescence imaging with CAV1, E-cadherin, β-actin, and DAPI (nuclei) (D). For RT-qPCT, data were normalised to fdAT2 organoids; mean ± SD, n = 3 independent repeats (*p<0.05, **p<0.01; unpaired t-test (two-tailed)). (E-G) AT1 differentiation of *SFTPC*-GFP-expressing fdAT2 organoids in a culture medium containing YAP signalling agonist, 10 μM LATS-IN-1, in the absence of Wnt agonists, in 3D organoid culture. Experimental scheme of AT1 differentiation (E), qRT-PCR analysis of AT1 markers *AQP5*, *CAV1*, and *AGER*, and an AT2 marker, *SFTPC* (F), and immunofluorescence imaging with CAV1, AGER, and DAPI (nuclei) (G). For RT-qPCR, data were normalised to fdAT2 organoids; mean ± SD, n=3 independent repeats (*p<0.05, **p<0.01; unpaired t-test (two-tailed)). (H) Explant culture of mouse precision-cut lung slices (PCLS) upon injecting *SFTPC*-GFP; *EF1a*-RFP human fdAT2 organoids. Organoids were dissociated and 2 x 10^5^ single cells mixed with 100 μl of 50% matrigel and directly injected into each lobe of the lungs. PCLS of 150 μm thickness were cultured for 1 week in 10% fetal bovine serum-containing medium. Three independent lines of fdAT2 organoids were used for the explant culture (16402, 16587, 16392). (I-K) Immunofluorescence images of PCLS showing engrafted and AT1 differentiated human cells. RFP+ cells were monitored by a combination of cell type marker antibodies against SFTPC (I,J), and CAV1 (I) or AGER (J). DAPI, nuclei. Scale bars, 50 μm. Quantitation (K) of AT1-lineage positive human cells in the explants, by measuring the proportion of AT2 (SFTPC) and AT1 markers (CAV1 and AGER) co-localising with RFP (n = 53, SFTPC/CAV1 cells; n = 56, SFTPC/AGER cells; 6 lung slices from 3 biological replicates).

Adult AT2 cells function as facultative progenitors of lung alveoli that can self-renew and differentiate into alveolar type 1 (AT1) cells to replenish AT1 cells upon injury^2,3^. We investigated whether the fdAT2 organoids have the capacity to differentiate into AT1-like cells. The fdAT2 organoids were treated with AT1 lineage-promoting conditions: 10% human serum medium on 2D for 2 weeks^10^, or 10 mM LATS-IN-1, an inhibitor of LATS1/2 kinases causing activated YAP signalling, in 3D for 1 week^23^ (Fig. 2A-G). In both conditions, the fdAT2 organoids showed upregulation of AT1 fate markers, such as *AQP5*, *CAV1*, and *AGER*, and downregulation of *SFTPC* and *SFTPC*-GFP (Fig. 2C,D,F,G). Next, we tested whether the fdAT2 organoids could differentiate in a more physiological environment. We dissected adult mouse lungs and injected them with single-cells isolated from *SFTPC*-GFP;*EF1a*-RFP-expressing human fdAT2 organoids, followed by precision-cut lung slice (PCLS) culture for 1 week. The RFP^+^ human cells were engrafted into the alveolar structure in the mouse PCLS and showed flattened nuclei, consistent with reports that AT1 cells have flattened nuclear shape^24^ (Fig. 2I-J; Supplementary Video 1). Scoring for the *SFTPC*-GFP reporter and AT1/2 fate markers confirmed that nearly 100% of the RFP^+^ cells co-expressed the AT1 lineage markers, CAV1 and AGER, and rarely expressed the *SFTPC*-GFP reporter or SFTPC protein (Fig. 2H-K). Taken together, these data show that the human fdAT2 organoids are competent to differentiate to the AT1 cell lineage *in vitro* and in the mouse lung environment *ex vivo*.

Our data indicate that the fdAT2 organoids self-renew, differentiate to AT1 cells and display a mature surfactant synthesis profile during prolonged passaging. They are amenable to lentiviral transduction and therefore represent a physiological system to study mechanisms of fundamental AT2 function and dysfunction in disease.

The commonest pathogenic variant of SFTPC, I73T, mislocalises to the plasma membrane and causes AT2 dysfunction via a toxic gain-of-function effect leading to ILD^6,7^. We were able to reproduce this cell surface phenotype in the fdAT2 organoids by viral transduction of HA-SFTPC variants (Fig. 3A,B). Ubiquitinated SFTPC is recognised by the ESCRT machinery and trafficked to late compartments. We previously showed that the disease-causing SFTPC-I73T mutant is no longer ubiquitinated, resulting in its relocalisation via recycling and saturated endocytosis of immature isoforms^8^ (Fig. 3C). To identify the ubiquitination machinery required for ESCRT recognition of SFTPC, we performed a targeted forward genetic screen (Fig. 3D). We predicted that depletion of key effectors of SFTPC ubiquitination would phenocopy the I73T variant by causing SFTPC accumulation at the plasma membrane. HeLa cells stably expressing GFP-SFTPC and Cas-9 were transduced with a subgenomic ubiquitome sgRNA library^25^, and cells with increased surface-localised pro-SFTPC when compared with untransduced controls were harvested at day 7 or 14 (Extended Data Fig. 3A,B). Transduced cells were also sorted for total SFTPC (GFP-high) to ensure that any cells accumulating C-terminally cleaved SFTPC at the cell surface (and thus missing the antibody epitope) were not overlooked (Extended Data Fig 3C,D).

The most enriched gRNA in d7 cell surface SFTPC-high cells targeted Itchy E3 Ubiquitin Protein Ligase (*ITCH)* (Fig. 3E; Extended Data Fig. 3A; Extended Data Table 4). Guide RNAs targeting the ESCRT machinery components Hepatocyte Growth Factor-Regulated Tyrosine Kinase Substrate (*HRS)* and VPS28 Subunit of ESCRT-I (*VPS28),* K63-chain specific Ubiquitin E2 Conjugating Enzyme E2 N (*UBE2N)* and the early endosome Rab5-specific GEF RAB Guanine Nucleotide Exchange Factor 1 (*RABGEF1)* were also enriched; supporting our previous data that SFTPC transits early endosomal compartments before K63-ubiquitination, recognition by the ESCRT complex and transit into MVBs. These hits largely overlapped with the GFP-defined sort which also included gRNAs targeting *ITCH* and ESCRT machinery (Extended Data Fig. 3C). Enrichment of gRNAs for sumoylation-related Ubiquitin Conjugating Enzyme E2 I (*UBE2I)*, Ubiquitin Like Modifier Activating Enzyme 2 (*UBA2)* and Protein Inhibitor Of Activated STAT 1 (*PIAS1)* was noted in both screens. *ITCH* remained highly enriched in the d14 sort, though other specific hits (e.g. *HRS*, *VSP28*, *UBE2N*) were lost likely due to their fundamental roles in trafficking and thus cellular toxicity when depleted (Extended Data Fig. 3A,B).

Initial validation of *ITCH*, SUMOylation components and positive controls *HRS* and *UBE2N* (Extended Data Fig. 4A) revealed that individual depletion resulted in marked cell surface GFP-SFTPC localisation (Extended Data Fig. 4B). Enrichment of full length SFTPC was more modest (Extended Data Fig 4C,D), suggesting that a proportion of surface-localised SFTPC is C-terminally cleaved. We used RT-qPCR to test the hypothesis that SUMOylation hits reflected changes in global transcription^26^, rather than SFTPC trafficking (Extended Data Fig. 4E) and based on these data SUMOylation was not investigated further.

ITCH is a HECT-type E3 ligase whose WW domains recognise cytosolic proline rich consensus sequences, typically PPxY which is present within the N-terminal tail of SFTPC^27^. Depletion of *ITCH* in clonal GFP-SFTPC expressing HeLa lines did not affect *SFTPC* transcription (Fig. 3F, I), but markedly increased plasma membrane resident SFTPC (Fig. 3G, H, J). Immunoblotting revealed an excess of full-length (*), partially cleaved (**) and fully C-terminally cleaved (***) species, suggesting that ITCH depletion inhibits trafficking at/beyond the location of the final C-terminal cleavage which is thought to be at the MVB limiting membrane^8,28^ (Fig. 3I). We confirmed that SFTPC resides largely in LAMP3^+^ (late) compartments in control cells. Following ITCH depletion, SFTPC relocalised to EEA1^+^ early endosomes and MICALL1^+^ recycling endosomes, consistent with failure of MVB entry and recycling to the plasma membrane (Fig. 3J). In rescue experiments, restoration of ITCH in knockout cells reversed SFTPC mislocalisation (Extended Data Fig. 5). These data suggest that ITCH is required for SFTPC trafficking and that ITCH depletion causes relocalisation of SFTPC to the plasma membrane, phenocopying SFTPC-I73T.

Having identified ITCH as an E3 ligase required for SFTPC maturation using HeLa cells, we wanted to determine whether ITCH also regulated endogenous SFTPC trafficking in primary AT2 cells. We therefore used CRISPRi to genetically deplete fdAT2 organoids of ITCH and UBE2N, as well as the HECT domain E3 ligase, NEDD4-2, which has been reported to play a role in SFTPC ubiquitination and maturation^29,30^. *NEDD4-2* was not isolated in our screen likely due to its relative lack of expression in HeLa (∼20% that of *ITCH;* www.ebi.ac.uk/gxa), but it is expressed in fdAT2 at a similar level to *ITCH* (Extended Data Fig. 2H).

CRISPRi-expressing fdAT2 organoids were transduced with a gRNA for each gene and silencing induced^31^ (Fig. 4A). Approximately 50% of organoids expressed both the CRISPRi and gRNA after 5 days of induction (Fig. 4B and Extended Data Fig. 6A,B). The toxicity of silencing ubiquitin ligases precluded sorting the double-positive population for RNA extraction, so the 40-60% overall gene depletion likely reflects complete gene depletion in dual-positive organoids (Fig. 4C). The knock-down effects reversed completely if organoids were recovered without silencing (-TMP/-dox) (Fig. 4D,E bottom panel ‘recovery’).

**Fig. 3:**
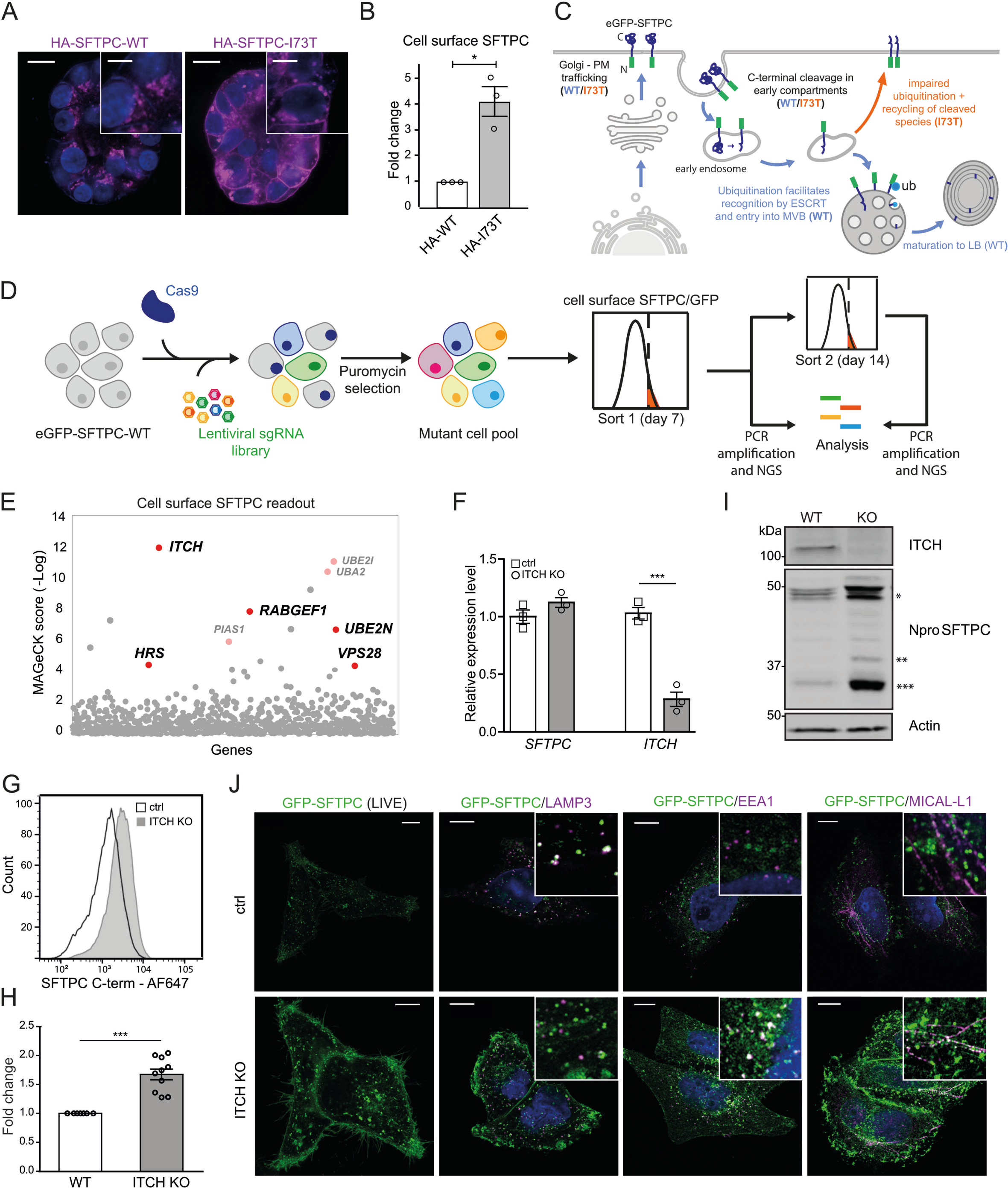
A forward genetic screen confirms the importance of ubiquitination in SFTPC trafficking and identifies the E3 ligase ITCH as required for SFTPC maturation. (A) Immunofluorescence of proSFTPC localisation in organoids expressing HA-SFTPC-WT or HA-SFTPC-I73T for 10 days. Scale bar 10 μm / 5 μm in zoomed inserts. (B) Quantification of cell surface SFTPC in organoids expressing HA-SFTPC-WT or HA-SFTPC-I73T as measured by SFTPC C-terminal antibody, expressed as fold change in mean fluorescence intensity; mean ± SEM, n=3 independent repeats (* = p<0.05, one-sample t-test). (C) Schematic of WT and pathogenic I73T variant SFTPC trafficking. After trafficking from early compartments to the plasma membrane, both WT and I73T are endocytosed and the C terminus cleaved in early endosomes. Onward trafficking into multivesicular bodies (MVBs) then lamellar bodies (LBs) is dependent upon further cleavage and ubiquitination; this fails in the I73T variant and results in recycling of partially cleaved isoforms to the plasma membrane. (D) Schematic of ubiquitome forward genetic screen strategy. HeLa cells stably expressing eGFP-SFTPC-WT and Cas9 were infected with a lentiviral ubiquitome sgRNA library consisting of 1,119 genes with 10 guides per gene. Transduced cells were selected with puromycin to generate a mutant cell pool. Day 7 post transduction, cells enriched for cell surface pro-SFTPC (as measured by a C-terminal antibody) or total GFP were isolated by FACS. These cells underwent a further enrichment sort at day 14. Genomic DNA was extracted from sorted cells and an unsorted control and genes with the highest gRNA representation identified by high-throughput sequencing. (E) MAGeCK score demonstrating relative enrichment of each gene for the day 7 cell surface SFTPC high population. (F) Relative expression of *SFTPC* and *ITCH* in control and ITCH knockout (KO) HeLa cells as measured by RT-qPCR and normalised to GAPDH; mean ± SEM, n=3 independent repeats (***=p<0.001, two-sided student’s t-test with non-equal variance). (G, H) Flow cytometry of cell surface SFTPC signal as measured by SFTPC C-terminal (C-term) antibody in control vs ITCH knockout lines and quantification of mean fluorescence intensity; mean ± SEM, n=10 independent repeats (*** = p<0.001, one-sample t-test). (I) Immunoblotting of lysates from control or ITCH knockout cells using an SFTPC N-terminal (Npro) and ITCH antibody (* = full length species (-/+ palmitoylation), ** = partially C-terminal cleaved intermediate, *** = fully C-terminally cleaved intermediate). (J) Live cell imaging and colocalisation of GFP-SFTPC with LAMP3, MICAL-L1 and EEA1 by immunofluorescence in control and ITCH knockout cells. Scale bar, 10 μm.

**Fig. 4:**
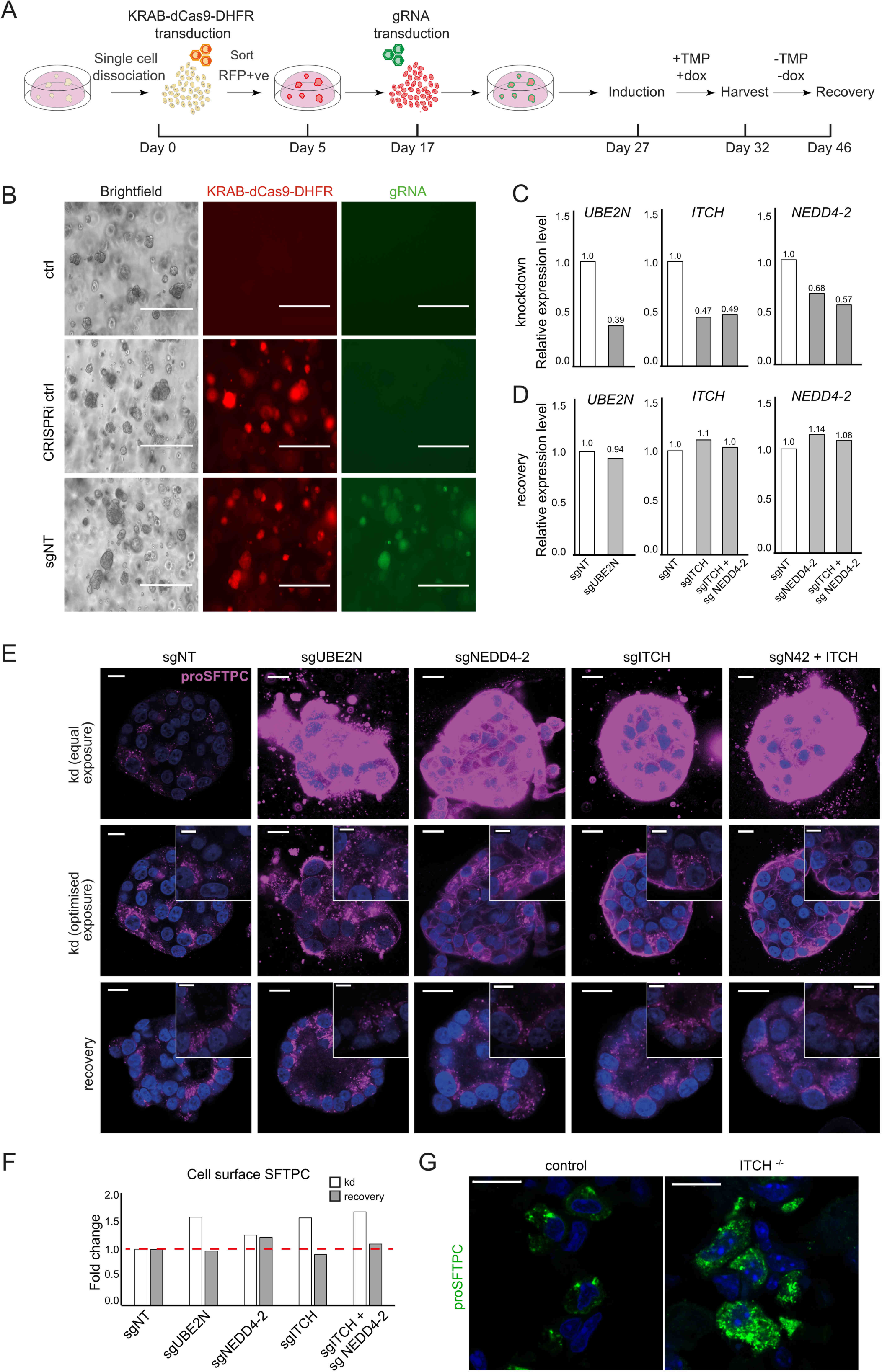
ITCH depletion alters SFTPC localisation in fetal lung-derived AT2 organoids. (A) Schematic of the inducible CRISPR interference (CRISPRi) system in fdAT2 organoids. CRISPRi organoid lines were generated by lentiviral transduction of AT2 cells with KRAB-dCas9-DHFR followed by sorting for RFP-positive cells. Cells were replated and expanded for 12 days before transduction with relevant sgRNA lentivirus. Approximately 10 days later the CRISPRi system was activated with doxycycline (dox) and trimethoprim (TMP). 5 days after induction, organoids were analysed by qPCR, flow cytometry and confocal microscopy. Dox and TMP were subsequently removed for 14 days to allow for recovery of gene expression. (B) Representative images showing morphology of control organoids and those transduced with KRAB-dCas9-DHFR (RFP) −/+ gRNA (GFP). Scale bar = 400 μm (C-D) Relative expression of *UBE2N, ITCH,* and *NEDD4-2* following CRISPRi induction (C) and 14-day recovery (D) (representative data from 3 (*ITCH/UBE2N*) or 1 (*NEDD4-2*) experiments, each performed in triplicate). Expression level was normalised to organoids expressing non-targeting control gRNAs. (E) proSFTPC localisation in CRISPRi-depleted (knockdown; kd) and recovered organoids by immunofluorescence (top panels: equal microscope settings illustrate accumulation of pro-SFTPC protein; middle panels: exposure altered to visualise subcellular localisation). Scale bar = 10 μm / 5 μm zoomed inserts. (F) Quantification of cell surface SFTPC signal measured by flow cytometry using SFTPC C-terminal antibody, expressed as fold change in mean fluorescence intensity. (G) Intracellular localisation of SFTPC in wild-type or Itch knockout mice.

Global inhibition of K63-ubiquitination and more targeted inhibition of ubiquitination (via *UBE2N* or *NEDD4-2* depletion respectively) resulted in massive accumulation of intracellular SFTPC with partial relocalisation to the plasma membrane (Fig. 4E top and middle rows). *ITCH* depletion resulted in a similar, but exaggerated, phenotype to that of *NEDD4-2* depletion, and dual-knockdown further exacerbated the intracellular and cell surface SFTPC accumulation. The cell surface relocalisation likely reflects both active recycling of C-terminally cleaved protein and an excess of full-length protein, which is detectable by flow cytometry (Fig. 4F and Extended Data Fig. 6C). Depletion of SFTPC ubiquitin ligases in human fdAT2 organoids results in mislocalisation of the endogenous SFTPC.

ITCH deficiency causes lung interstitial inflammatory infiltrates in humans and mouse models^32,33^ thought to result from multisystem autoimmune disease. We predicted an additional alveolar epithelial phenotype if ITCH is important for SFTPC maturation *in vivo.* Staining for pro-SFTPC revealed intracellular accumulation and an altered distribution in *Itch* deficient (*Itch^a18H/a18H^*) mouse alveolar epithelium. Numerous, smaller pro-SFTPC puncta are consistent with failure of trafficking to late compartments, seen as large puncta in the wild-type mice (Fig. 4G).

We conclude that intracellular trafficking of SFTPC requires K63 ubiquitination mediated by the HECT domain E3 ligases including ITCH and, to a lesser extent, NEDD4-2. Inhibiting ubiquitination phenocopies the redistribution of the pathogenic SFTPC-I73T variant, reinforcing the importance of this post-translational modification in maintaining AT2 health.

We have derived an expandable AT2 organoid model from fetal lung tip progenitor cells and demonstrated its use in investigating mechanisms of AT2 biology relevant to disease. FdAT2 organoids acquire features of mature adult AT2 cells, including lamellar body formation, mature surfactant protein secretion, and the ability to differentiate to AT1 cells (Figs. 1,2). In contrast to human adult lung-derived AT2 organoids^9–11^, the fdAT2 organoids readily expand, can be passaged multiple times and cryopreserved without compromising their identity. Their transcriptome is similar to that of PSC-iAT2 (Fig. 1G), but they can be derived in a more straightforward and timely fashion. They are also amenable to genetic manipulation, enabling us to investigate key SFTPC trafficking effectors using CRISPRi (Figs. 3,4).

Complete trafficking and maturation of SFTPC is required for AT2 health; misfolding variants are retained in early compartments and cause ER stress^34,35^ whereas mistrafficking isoforms, exemplified by SFTPC-I73T, mislocalise due to a trafficking block and failure of ubiquitination^8,35,36^. Both mutation types result in AT2 cell dysfunction and heritable forms of interstitial lung disease. These phenotypes have been confirmed in animal models^37–39^, but mechanistic work on AT2 dysfunction in physiological human *ex vivo* models has been highly challenging. We combined a forward genetic screen with genetic manipulation of fdAT2 organoids to investigate intracellular SFTPC trafficking via the plasma membrane (Fig. 1D). We confirm that ubiquitination is required for normal SFTPC trafficking and identify the E3 ligase ITCH as a novel effector of SFTPC processing. Although there can be redundancy between family members, the observation that individual depletion of either ITCH or NEDD4-2 (expressed at similar levels in human AT2 cells) yields a phenotype close to that of dual depletion (Fig. 4E) suggests they may play complementary roles. We further confirmed altered SFTPC handling in *Itch* deficient mice (fig 4G).

FdAT2 organoids are an attractive genetic model for understanding fundamental mechanisms of AT2 cell biology in health, and modelling inherited disorders and environmental insults which perturb their function. They also represent a useful cell source to investigate cellular and molecular mechanisms of AT2 to AT1 cell lineage differentiation during human lung development and repair.

## Acknowledgements

We would like to acknowledge the imaging facilities at the Gurdon Institute and Cambridge Institute for Medical Research and the flow cytometry facility at the Cambridge Institute for Medical Research (CIMR).

KL is supported by the Basic Science Research Program through the National Research Foundation of Korea (NRF) funded by the Ministry of Education (2018R1A6A3A03012122). ENR and JAD are supported by an MRC Clinician Scientist Fellowship (MR/S005552/1). DS is supported by a Wellcome Trust PhD studentship (109146/Z/15/Z) and the Department of Pathology, University of Cambridge. JRE is supported by a Sir Henry Dale Fellowship jointly funded by the Wellcome Trust and the Royal Society (216370/Z/19/Z). LEM was supported in part by the National Institute of General Medical Sciences of the National Institutes of Health under Award Number P20GM103499. J-HL and JHB are supported by a Wellcome Senior Research Fellowship (221857/Z/20/Z) and the Suh Kyungbae Foundation (SUHF-20010033). SJM is supported by the MRC (MCMB MR/V028669/1 and MR/R009120/1), EPSRC (EP/R03558X/1), Cambridge Biomedical Research Centre (BRC-1215-20014); Asthma+Lung UK (ALUK), and the Victor Philip Dahdaleh Foundation. ELR is supported by the Medical Research Council (MR/P009581/1). PJL and DJHVB are supported by a Wellcome Trust Principal Research Fellowship (210688/Z/18/Z) and an MRC project grant (MR/V011561/1).

We acknowledge core funding to the Gurdon Institute from the Wellcome Trust (203144/Z/16/Z) and CRUK (C6946/A24843). This research was also supported by the CIMR Flow Cytometry Core Facility.

## Author contributions

Conceptualisation, KL, ENR, PJL, SJM, ELR, JAD; Methodology, Investigation, and Validation, KL, ENR, DS, DJHVB, JRE, LEM, JAD; Resources, LEM, PJL; Software and Formal Analysis, KL, ENR, DJHVB; Writing – Original Draft KL, ENR; Writing – Review & Editing, JHL, PJL, SJM, ELR, JAD; Funding Acquisition and Supervision: JHL, PJL, SJM, ELR, JAD.

---

**Extended Data Fig. 1.**
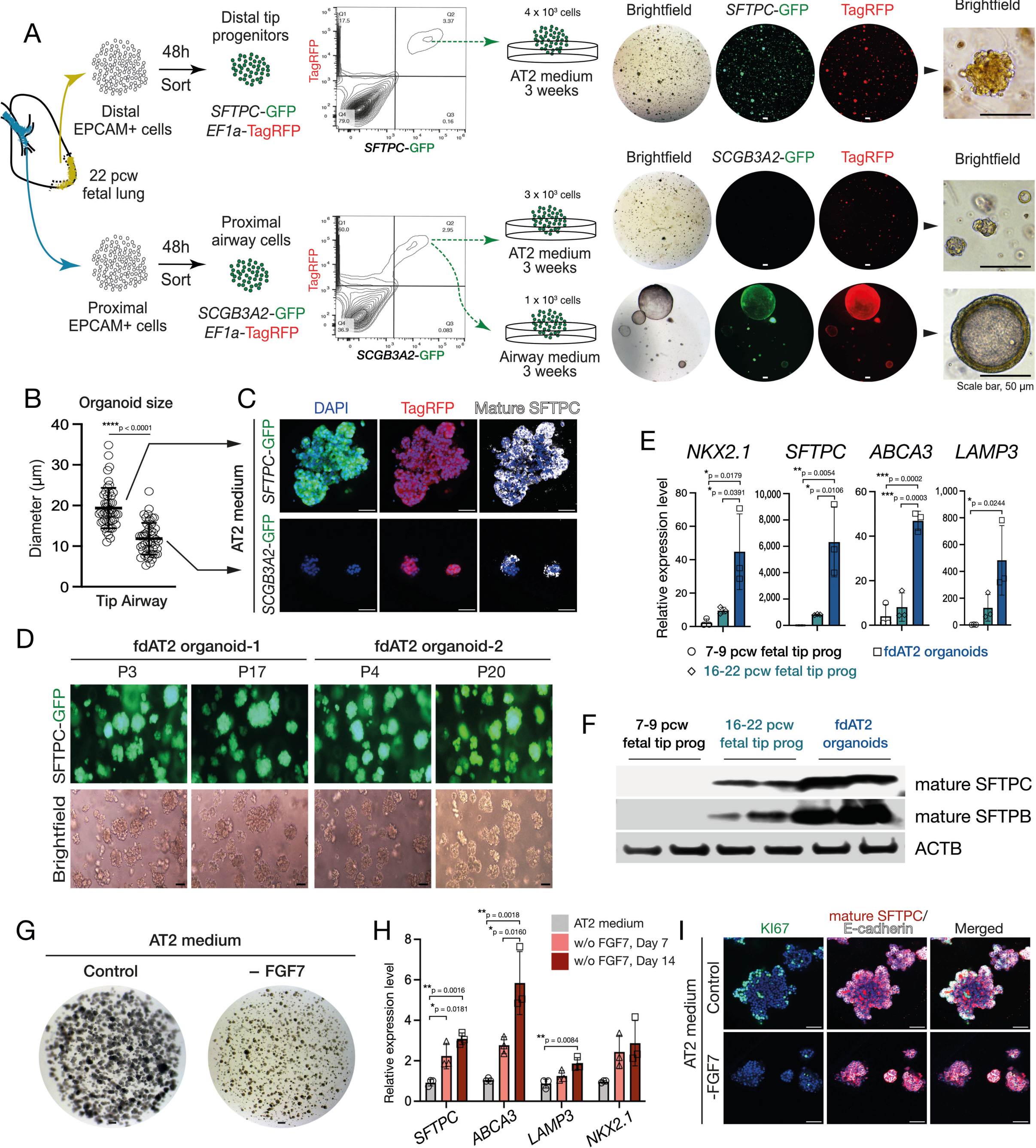
Characterization of AT2 organoids. (A) Derivation and establishment of fdAT2 organoids from lung tip progenitor cells (upper panel), or proximal airway progenitors (lower panel) of human fetal lungs at 22 pcw. Upper panel: The isolated tip progenitor cells were immediately transduced and selected based on *SFTPC*-GFP and EF1a-TagRFP after 48 h of transduction; *SFTPC*-GFP and EF1a-TagRFP reporter positive cells were efficiently expanded into fdAT2 organoids when grown in AT2 medium for 3 weeks. Lower panel: Proximal airway cells were immediately transduced with *SCGB3A2*-GFP, EF1a-TagRFP reporter lentivirus and the airway progenitor cells were selectively isolated by *SCGB3A2*-GFP after 48h of transduction; *SCGB3A2*-GFP, *EF1a*-TagRFP reporter positive cells and expanded into small AT2-like organoids when grown in AT2 medium for 3 weeks, but efficiently formed airway organoids when grown in airway medium for the same period. Scale bar, 50 μm (B) Size of organoids expanded from tip progenitors (*SFTPC*-GFP^+^) or proximal airway progenitors (*SCGB3A2*-GFP^+^) in AT2 medium was measured; mean ± SD, n=50 (****p<0.0001; unpaired t-test (two-tailed)). (C) Expression of mature SFTPC protein in organoids expanded from tip progenitors or airway progenitors in AT2 medium. DAPI, nuclei. Scale bar, 50 μm. (D) Cultured fdAT2 organoids at early and late passages, stably expressing *SFTPC* promoter-driven GFP (*SFTPC*-GFP). Two independent lines of AT2 organoids at P3 and P17, and P4 and P20. Scale bars, 50 μm. (E) qRT-PCR analysis of alveolar type 2 cell lineage markers, *NKX2*.1, *SFTPC*, *ABCA3*, and *LAMP3*, in 7-9 pcw and 16-22 pcw tip progenitor organoids, and fdAT2 organoids at P12, P20, and P21. Data were normalised to 7-9 pcw tip organoids; mean ± SD, n=3 independent repeats (*p<0.05, **p<0.01, ***p<0.001; One-way ANOVA with Tukey multiple comparison post-test). (F) Immunoblot of mature forms of SFTPC and SFTPB in human fetal lung tip progenitor-derived organoids at 7-9 pcw and 16-22 pcw, respectively, and the fdAT2 organoids. Two independent replicates were used. (G - I) FdAT2 organoids were cultured in the AT2 medium for 2 weeks, in the presence (control) or absence of FGF7 (-FGF7) and AT2 lineage markers were measured by qRT-PCR (H) after 7 and 14 days of culture. Data were normalised to AT2 organoids cultured in the AT2 medium containing FGF7 (control); mean ± SD, n=3 independent repeats (*p < 0.05, **p < 0.01, ***p < 0.001; one-way ANOVA with Tukey multiple comparison post-test). (I) Mature SFTPC protein expression was visualised with a proliferation marker, KI67, and E-cadherin by immunofluorescence staining, at 14 days of culture. DAPI, nuclei. Scale bar, 50 μm.

**Extended Data Fig. 2.**
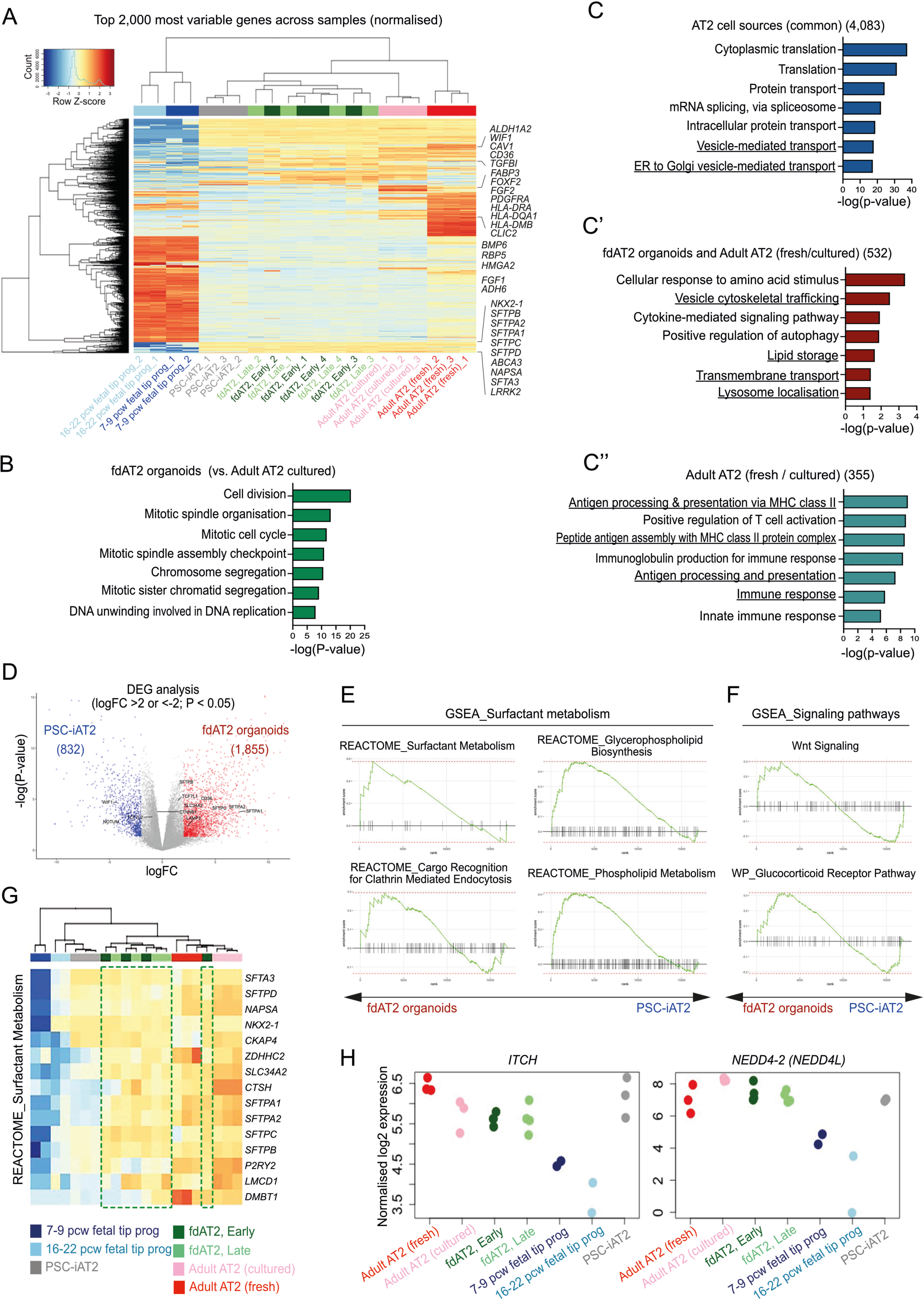
Comparative transcriptomic analysis of fetal-derived AT2 organoids with other AT2 sources. (A) Heatmap analysis of the top 2,000 most variable genes across all samples. (B) GO analysis of genes highly enriched in the fdAT2 organoids compared to the cultured adult AT2 cells. (C, blue) GO analysis of DEGs shared between AT2 cells of different origin, including fdAT2 organoids, PSC-iAT2, and cultured and freshly isolated adult AT2 cells; related to Fig. 1H. 4,083 genes commonly shared by all AT2 fate cell types. (C’, red) genes shared by fdAT2 organoids and cultured and/or freshly isolated adult AT2 cells (C’’; cyan) genes shared by cultured and freshly isolated adult AT2 cells. (D) Volcano plot describing the direct comparison of fdAT2 organoids and PSC-iAT2. 1,855 and 832 genes were differentially enriched in AT2 organoids and PSC-iAT2, respectively (logFC > 2, *P*-value < 0.05; related to Extended Data Table 3). (E and F) Gene set enrichment analysis (GSEA) of surfactant metabolism and signalling pathway-associated gene sets between fdAT2 organoids and PSC-iAT2. (G) Heatmap of a gene set associated with surfactant metabolism from REACTOME. Green box, fdAT2 organoids. (H) Relative expression of E3 ligases *ITCH* and *NEDD4-2* in AT2 cells and tip progenitor organoids. .

**Extended Data Fig. 3.**
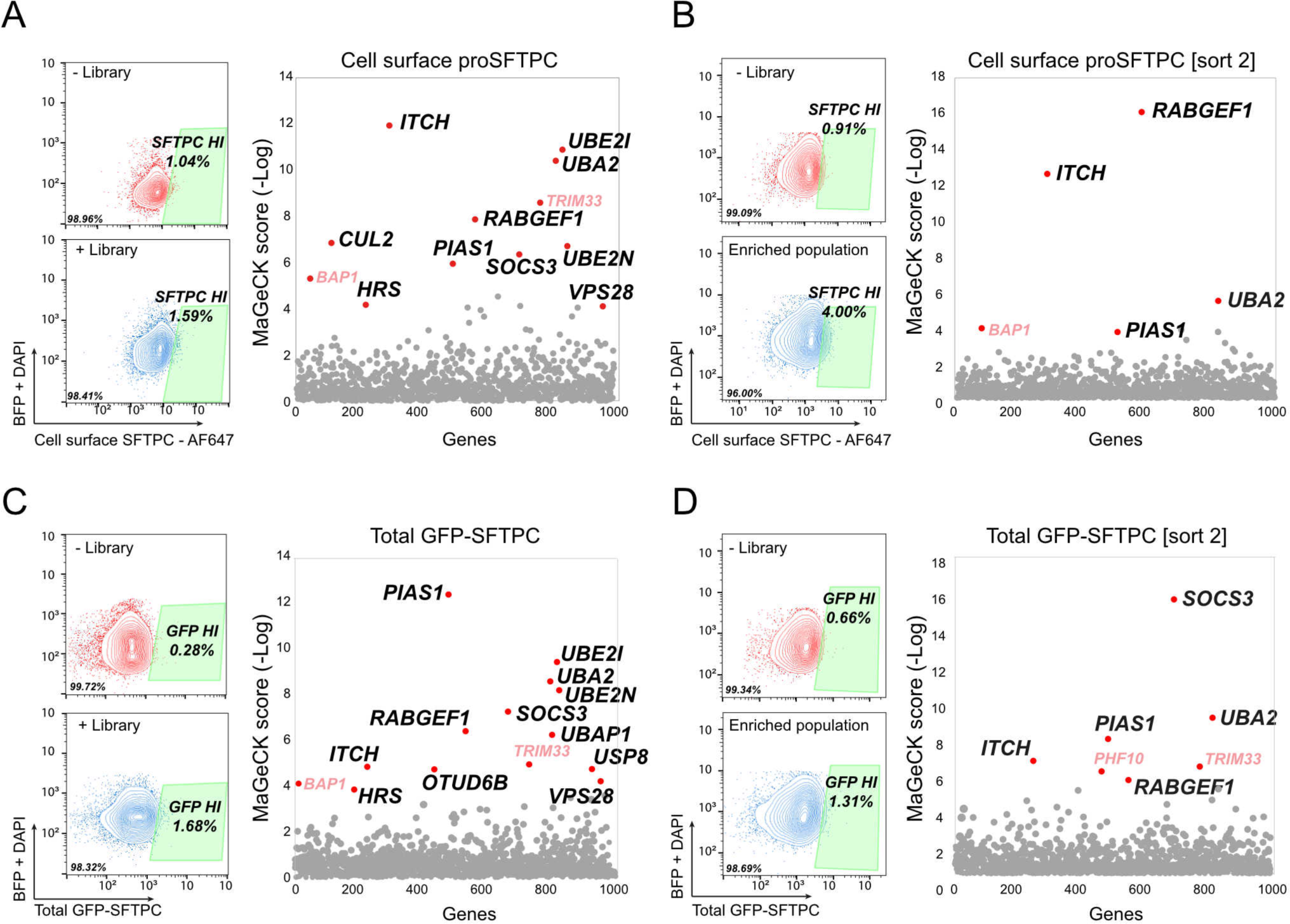
A forward genetic screen identifies candidate proteins involved in SFTPC processing and trafficking. (A-B) Flow cytometry gating strategy and MAGeCK relative enrichment scores for genes whose depletion results in increased cell surface SFTPC (A, day 7 and B, day 14) or increased total eGFP-SFTPC (C, day 7 and D, day 14) post-transduction with ubiquitome sgRNA library. Genes highlighted red (BAP1,TRIM33 and PHF10) are commonly enriched but non-specific transcription-related hits from forward genetic screens.

**Extended Data Fig. 4.**
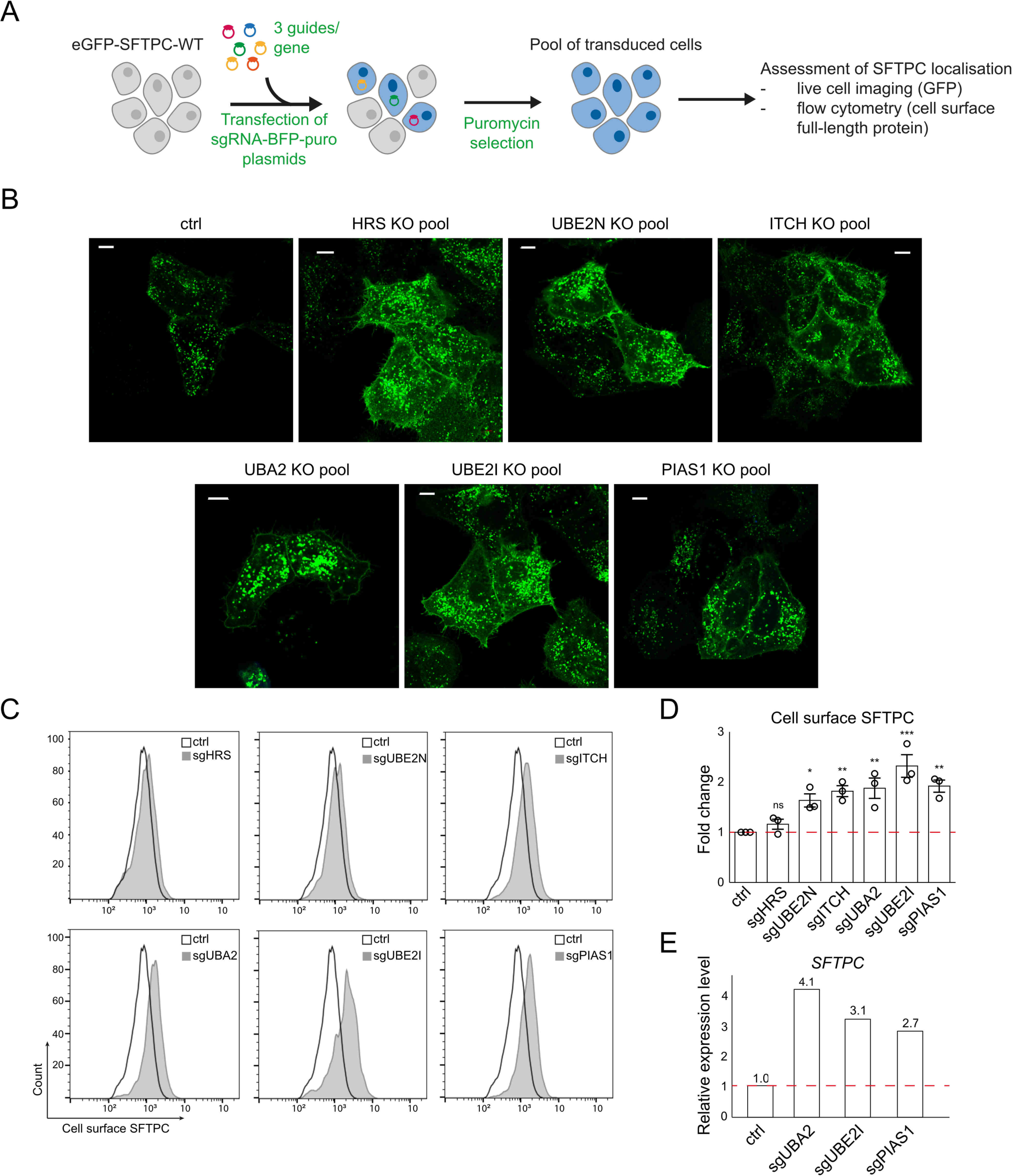
Initial validation of screen hits. (A) Schematic of screen hit validation strategy. HeLa cells stably expressing GFP-SFTPC-WT and Cas9 were transfected with plasmids containing sgRNAs against specific genes for hit validation. Three guides for each gene were pooled for transfection. Cells underwent puromycin selection for 48 hours to select for transfected cells. (B-C) Knockout pools assessed for cell surface SFTPC enrichment by live cell confocal microscopy (B) and flow cytometry (C). Scale bar = 10 μm. (D) Quantification of full-length cell surface SFTPC as measured by C-terminal antibody and expressed as mean fluorescence intensity. Mean ± SEM, n=3 independent repeats (*p < 0.05, **p < 0.01,***p<0.001; paired two-tailed Student’s *t*-test). (E) Relative expression of *SFTPC* mRNA in GFP-SFTPC-Cas9 control cells and UBE2I, UBA2, and PIAS1 knockout pools (representative result of 3 independent repeats).

**Extended Data Fig. 5.**
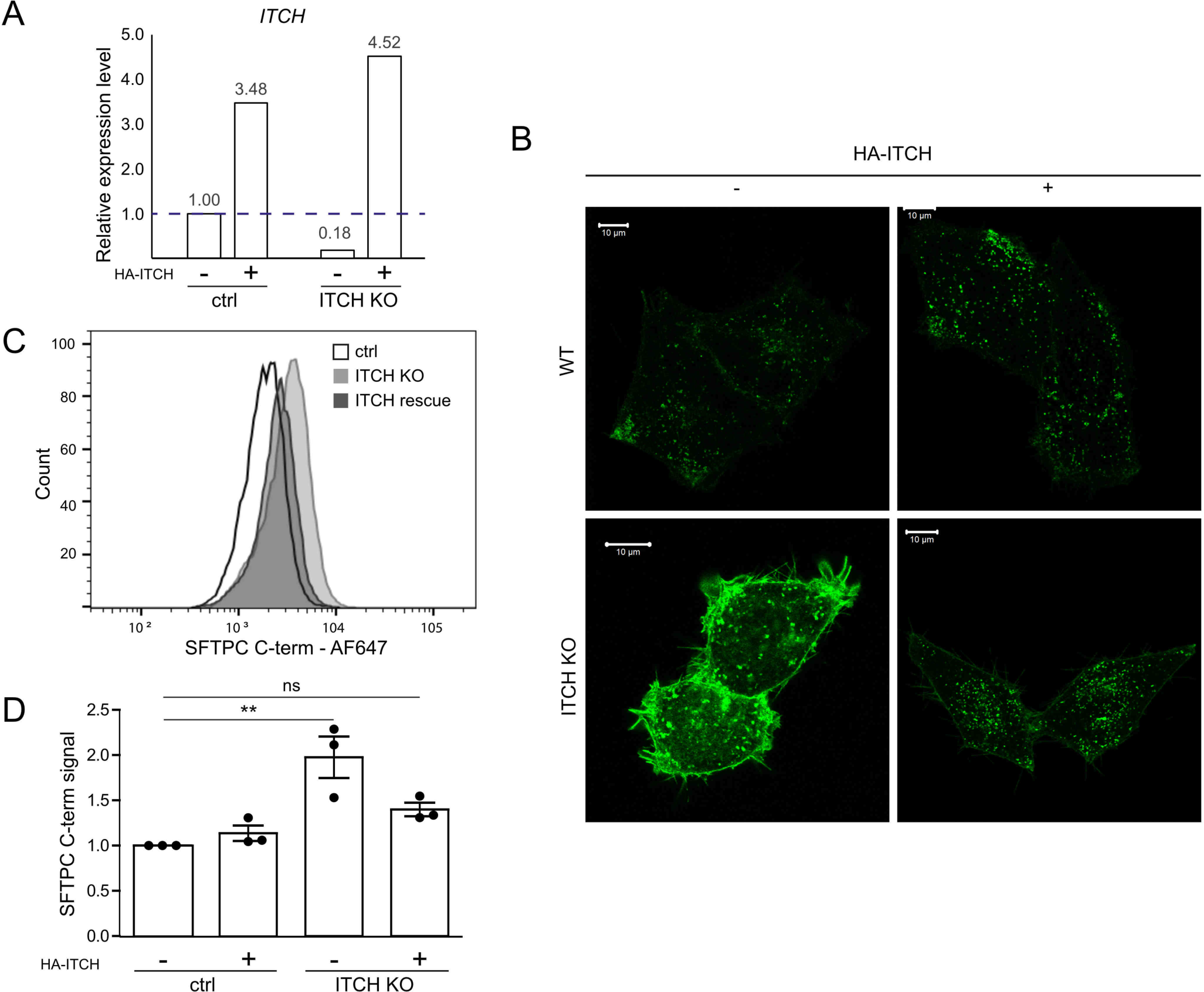
Restoration of ITCH expression in knockout cells reverses SFTPC mislocalisation. Control and ITCH knockout HeLa cells transfected with *HA-ITCH* were assessed for relative *ITCH* mRNA expression (A), GFP-SFTPC localisation (B) and cell surface full-length SFTPC protein by flow cytometry (C&D). Scale bar = 10 μm. Mean ± SEM, n=3 independent repeats (**=p<0.005, one-way ANOVA with *post-hoc* Tukey tests).

**Extended Data Fig. 6.**
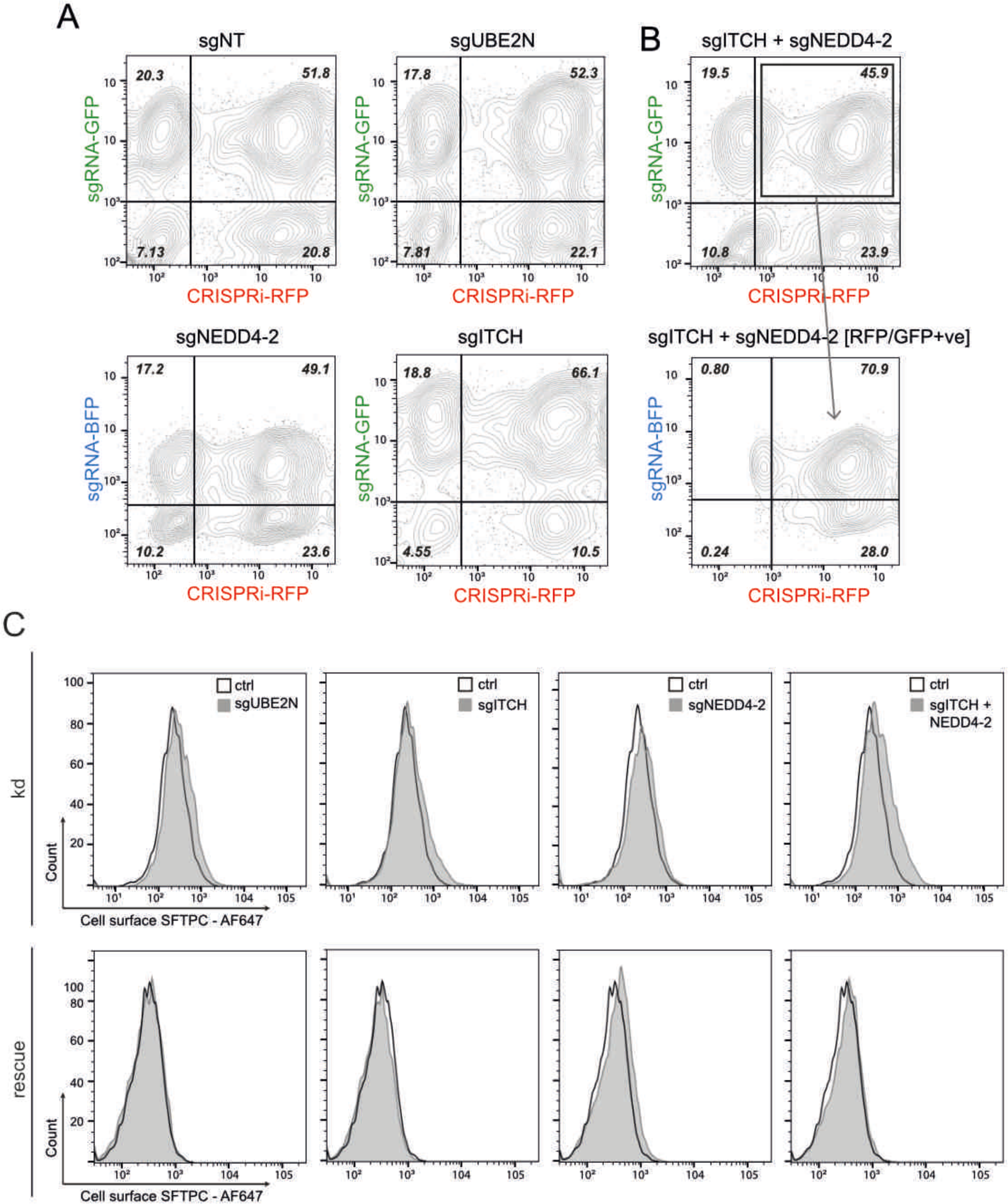
Flow cytometry gating strategy and cell-surface SFTPC measured during CRISPRi knockdown and recovery. (A) Flow cytometry dot plots of sgRNA positive (y-axis) and CRISPRi-RFP (x-axis) positive populations in organoids transduced with sg non-targeting (NT), sgUBE2N, sgITCH, or sgNEDD4-2 at cell harvesting (5 days post induction of CRISPRi system). (B) (top panel) Flow cytometry dot plots of sgRNA positive (y-axis) and CRISPRi-RFP (x-axis) positive populations for cells transduced with both sgITCH and sgNEDD4-2 (5 days post induction of CRISPRi system). The population was first gated for GFP and RFP positive cells. The resulting population (outlined) was gated for BFP positive cells (bottom panel). (C) Flow cytometry of cell surface SFTPC as measured by C-terminal antibody (detecting full length protein) during knockdown (kd) and following rescue when compared with organoids expressing non-targeting control gRNAs.

**Extended Data Table 1. Multiple comparison of transcriptomes of alveolar type 2 cells of different origins.** Transcriptional comparison between fdAT2 organoids with other AT2 cells.

**Extended Data Table 2. Transcriptomic comparison of AT2 cells compared to those of fetal 16-22 pcw tip progenitor organoids.** The genes for each cell type were differentially extracted from those of fetal late tip progenitor organoids (logFC > 2, *P*-value < 0.05).

**Extended Data Table 3. Direct comparison of transcriptomes of fetal-derived AT2 organoids and PSC-iAT2.** Transcriptional comparison between fdAT2 organoids with PSC-iAT2.

**Extended Data Table 4**. Full list of enriched genes in ubiquitome forward genetic screens at day 7 and day 14.

## Methods and protocols

Details of primary antibodies, expression plasmids, primers, guide RNA, and sample information are included in supplementary table 1.

## Material Availability

Human organoid lines used in the study are available from the Dr Emma L Rawlins with a completed Materials Transfer Agreement.

## Mouse tissue

6-10 week-old C57BL/6 mice were used for the ex vivo AT1 differentiation experiments. All procedures were approved by the University of Cambridge Animal Welfare and Ethical Review Body and carried out under a UK Home Office License (PPL: PP3176550) in accordance with the Animals (Scientific Procedures) Act 1986. Mice were bred and maintained under specific-pathogen-free conditions at the Gurdon Institute of the University of Cambridge.

Twelve-week old animals homozygous for a null allele of *Itch* (*Itch^a18H/a18H^*, B6.C3H(101)-In(2a;Itch)18H/LmatMmjax MMRRC stock #65285) have been previously described (Hustad et al., 1995; Cattanach BM et al, 1987). For this line, the *a^18H^*allele was backcrossed to C57BL/6J for 27 generations. Therefore, age- and gender-matched C57BL/6J mice were used as controls (*Itch^+/+^*) in the indicated experiments. All mice were cared for in accordance with the National Institute of Health’s Guide for the Care and Use of Laboratory Animals (8th ed., Washington, DC: 2011) and the University of South Carolina’s Institutional Animal Care and Use Committee approved all experimental protocols.

## Human embryonic and fetal lung tissues

Human embryonic and fetal lung tissue was provided from terminations of pregnancy from Cambridge University Hospitals NHS Foundation Trust under permission from NHS Research Ethical Committee (96/085) and the MRC/Wellcome Trust Human Developmental Biology Resource (London and Newcastle, University College London (UCL) site REC reference: 18/LO/0822; Project 200591; www.hdbr.org). Sample age ranged from 7-9^1^ and from 16-22 weeks of gestation (post-conception weeks; pcw). Sample gestation was determined by external physical appearance and measurements. Samples had no known genetic abnormalities. Sample gender was unknown at the time of collection and was not determined. All collected samples were included in the study.

## Derivation and *in vitro* culture of human alveolar type 2-like organoids

Distal edges of 16∼22 pcw human fetal lung tissue, typically measuring 0.5 cm x 1∼3 cm, were sectioned and fragmented into smaller pieces. Fragments were dissociated with 0.125 mg/ml Collagenase (Merck, C9891), 1 U/ml Dispase (Thermo Fisher Scientific, 17105041), and 0.1 U/μl DNase (Merck, D4527) in a rotating incubator for 1 hr at 37°C. After rinsing in washing buffer containing 2% FBS in cold PBS the cells were filtered through a 100 μm strainer. Cells were treated with RBC lysis buffer (BioLegend, 420301) at room temperature for 5 min, rinsed and incubated with EpCAM (CD326) microbeads (Miltenyi Biotec; 130-061-101) to isolate EpCAM+ epithelial cells containing mostly tip epithelial cells. These cells were embedded in matrigel (Corning, 356231) and cultured in 24-well plates in alveolar type 2 differentiation medium (AT2 medium): Advanced DMEM/F12 supplemented with 1x GlutaMax, 1 mM HEPES and Penicillin/Streptomycin, 1X B27 supplement (without Vitamin A), 1X N2 supplement, 1.25 mM N-acetylcysteine, 50 nM Dexamethasone (Merck, D4902), 0.1 mM 8-Bromoadenosine 3’5’-cyclic monophosphate (cAMP; Merck, B5386), 0.1 mM 3-Isobutyl-1-methylxanthine (IBMX; Merck, 15679), 50 μM DAPT (Merck, D5942), 100 ng/ml recombinant human FGF7 (PeproTech, 100-19), 3 μM CHIR99021 (Stem Cell Institute, University of Cambridge), 10 μM A83-01 (Tocris, 2939), and 10 μM Y-27632 (Merck, 688000). Medium was replaced every 2 days and cultures maintained for 2 weeks until the initial AT2 organoid colonies formed in the matrigel droplet. Organoids were typically passaged at a 1 to 3 ratio weekly by gently breaking into small fragments.

For SFTPC-GFP, or SCGB3A2-GFP^2^ reporter cell isolation, the isolated tip progenitor cells or proximal airway cells were immediately transduced and selected based on *SFTPC*-GFP or *SCGB3A2*-GFP with *EF1a*-TagRFP reporter expression by flow cytometry, after 48 h of transduction and cultured for 3 weeks in the AT2 medium. As a control, proximal airway progenitors were immediately selected based on *SCGB3A2*-GFP after 48 h of transduction, and cultured in airway medium: Advanced DMEM/F12 medium supplemented with 1X B27, 1X N2, 1.25 mM N-acetylcysteine, 100 ng/mL FGF10, 100 ng/mL FGF7, 50 nM Dexamethasone, 0.1 mM cAMP, 0.1 mM IBMX, and 10 μM Y-27632.

## Immortalised cell culture and cell line derivation

HeLa cells were cultured in DMEM (Sigma-Aldrich, D6429) + 10% fetal bovine serum (FBS) (Sigma-Aldrich). Plasmid DNA was introduced using liposomal transfection with FuGene 6 (Promega) or lipofectamine LTX (ThermoFisher).

Clonal GFP-SFTPC-expressing HeLa cell lines were derived as previously described^3^. A pool stably expressing Cas9 was generated by lentiviral transduction of the pHRSIN-Cas9 vector and selection of transduced cells using hygromycin (2 μg/ml; Invitrogen 10687010).

For initial CRISPR screen validation, gene-specific deplete pools were obtained by transducing HeLas with pKLV-pU6-esgRNA(modified BbsI)-pPGK-Puro2ABFP containing 3 gRNAs per gene and transduced cells selected with puromycin (1 μg/ml; Alfa Aesar J61278). *ITCH* depleted clonal lines were subsequently derived by single cell sorting into 96 well plates before expansion and screening by RT-qPCR. Experiments were performed on a minimum of 3 clonal lines derived from different gRNAs to ensure findings did not reflect off-target effects.

For ITCH rescue experiments, 4 silent mutations were introduced into the *ITCH* cDNA sequence in the region of the gRNA by site-directed mutagenesis (GAA CGG CGG GTT GAC AAC ATG -> GAG CGG CGG GTA GAT AAT ATG) before transfection to ensure the integrated CRISPR-Cas9 in the ITCH knockout cells did not also edit the transiently transfected plasmid.

To determine the efficacy of candidate CRISPRi gRNAs, HeLas were transduced with pLenti-tetON-KRAB-dCas9-DHFR-EF1a-TagRFP-2A-tet3G and sorted by FACS. Expression was induced with TMP and dox for 5 days before cells harvested for RNA and RT-qPCR.

## Alveolar type 1 (AT1) lineage differentiation of AT2 organoids *in vitro* and *ex vivo*

For *in vitro* AT1 differentiation, AT2 organoids were dissociated into single cells and 1 x 10^5^ cells were replated on matrigel-coated 12-well dishes and cultured either 1) in an AT1-promoting medium^4^ containing 10% human serum for 2 weeks or 2) in medium^5^ containing the LATS1 and 2 inhibitor LATS-IN-1 (10 mM, Cambridge Bioscience, CAY36623) and in the absence of CHIR99021 for a week. Cells were analysed by RT-qPCR and immunofluorescence.

AT1 lineage differentiation of AT2 organoids was performed in an *ex vivo* mouse lung explant. Prior to the *ex vivo* culture of precision-cut lung slices (PCLS)^6^, AT2 organoids expressing *SFTPC*-GFP+ and TagRFP+ were dissociated into single cells, 2 x 10^5^ cells mixed with 100 μl 50% matrigel and directly injected into each lung lobe using an 18G needle. Mouse lung lobes were sectioned into 150 μm PCLS using a Leica VT1200s vibratome. Following *ex vivo* culture in DMEM/F12 medium supplemented with 2% v/v penicillin-streptomycin, 10% FBS and 1 mM cAMP for one week, the PCLS were subjected to immunofluorescence analysis.

## Lentivirus production and viral transduction

Lentivirus was produced using HEK293T cells. For pHRSIN and pKLV vectors (for the forward genetic screen), 5 x 10^5^ cells were plated in one well of a 6-well plate 24 hr before transfection with 0.67μg pCMVR8.91 (Gag-Pol), 0.33μg pMD2G (VSV-G) and 1μg pHRSIN/pKLV (with gene of interest) and media was changed the next day. Virus was collected after 48 hr and filtered through a 0.45μm filter. For pLenti vectors for the CRISPRi experiments and generating SFTPC-GFP reporter line, 2 x 10^5^ / 8 x 10^5^ cells (<10kb / >10kb plasmid, respectively) were plated into a 10cm dish 24 hr before transduction with 5/10μg plasmid of interest, 3/6μg pPAX2, 2/3μg pMD2.G and 2/3μg pAdvantage and media was changed the next day. Virus was collected after 48 hr and filtered through a 0.45 μm filter before being concentrated with Lenti-X (Takara bio) (3:1 supernatant to lenti-X) overnight at 4°C, spun at 1,500 g for 45 min in a cold centrifuge and the pellet resuspended such that virus was 100X concentrated.

Transduction of HeLa cells was typically achieved by adding 500 μl viral supernatant to 2 x 10^5^ cells in a 6-well plate, centrifuging at 600 x g for 1 hr and incubating overnight before refreshing the media. Antibiotic selection was added 48 hr after transduction.

## Genetic manipulation of organoids

To create transduced organoid lines, single-cell dissociated AT2 organoids were infected with lentivirus overnight at 37°C then embedded into matrigel and cultured in the medium for another 48-72 before fluorescence sorting to enrich for transduced cells. For the reporter system, pHAGE plasmids were modified by insertion of EGFP or EF1a-promoter TagRFP (EF1a-TagRFP) cassettes into the human *SFTPC* promoter (2.2 kb; chr8:22,433,535-22,435,769). For the HA-SFTPC lines, HA-SFTPC-WT or I73T were cloned into pLenti-tetON-EF1a-tagRFP-2A-tet3G by Gibson assembly.

For CRISPRi, the organoids were initially transduced with doxycycline-inducible lentivirus^7^ containing dCAS9 protein fused with 5’ KRAB and 3’ DHFR, and EF1a-TagRFP vector Post-sorting, organoids were expanded before being transduced with lentivirus harbouring U6 promoter driven guide RNAs and EF1a-EGFP-CAAX, with the relevant guides^8^. For activation of the CRIPSRi system, doxycycline (2 μg/ml, Merck, D9891) and trimethoprim (10 nmol/l, Merck, 92131) were added to the medium for 5 days. For rescue experiments, doxycycline and trimethoprim were removed from the media for 14 days and organoids passaged where necessary before harvesting. CRISPRi experiments were carried out in two biologically independent lines.

## CRISPR screen

The forward genetic screen was undertaken using a ubiquitome gRNA library consisting of approximately 1,119 genes with 10 guides per gene^9^. The Cas9 activity of the HeLa-Cas9 line was confirmed by transducing with pKLV encoding β-2 microglobulin gRNA and assessing loss of MHC class I from the plasma membrane by flow cytometry (with >80% loss considered acceptable). The amount of virus required for an MOI of 0.3 was determined by transducing 1×10^6^ cells with varying amounts (25-400 μl) lentivirus and assessing BFP positivity by flow cytometry at 72 hr post transduction. For the screen, 20 million cells were transduced to ensure 500-fold coverage and puromycin selection commenced at 48 hr post transduction. After 5 days of selection, cells were harvested with 10mM EDTA and stained using the SFTPC BRICHOS domain antibody. Cells most highly enriched for BRICHOS signal or GFP (approx. 1%) were collected; half were harvested for genomic DNA extraction and half kept in culture for a further 7 days. This population underwent a further sort and the BRICHOS or GFP enriched population harvested. Genomic DNA was extracted using a Qiagen Puregene Core Kit (#1042601). Lentiviral gRNA inserts were amplified in a two-step PCR reaction as previously described (Menzies et al, 2018), cleaned with AMPure XP magnetic beads (Beckman Coulter A63881) and sequenced by MiniSeq (Illumina).

## Flow cytometry

HeLa cells were detached using 10 mM EDTA and organoids dissociated to single cells before being washed with PBS and centrifuged at 500 g for 3 min to pellet. Non-specific staining was blocked with 10% FBS in PBS for 30 min then cells pelleted by centrifugation and incubated with primary antibody on ice for 30 min. Following two rounds of washing with PBS by centrifugation, cells were incubated with AF647-conjugated secondary antibodies for 30 min on ice, washed twice and filtered through 50 μm filters. Samples were analysed on a Fortessa (BD bioscience) flow cytometer and further analysis undertaken using FlowJo software (10.0.0).

Flow cytometric sorting of single-cell dissociated organoids transduced with fluorescent markers was performed using a sorter (SH800S or BD FACSMelody) and analysed using FlowJo.

## Immunoblotting

For immunoblotting, cells were subjected to triton lysis, SDS-PAGE electrophoresis, and immunoblotting as previously described^10^. Membranes were incubated overnight with primary antibody at 4°C before washing and incubating with secondary antibody (1:20,000 IRDye® conjugated, various) for 1 hour at room temperature. Membranes were visualised using a Li-Cor Odyssey imaging system.

## Immunofluorescence staining

For immunostaining of Hela cells, cells grown on coverslips were fixed with 4% PFA for 30 min, blocked with 10% FBS for 30 min then permeabilized for 30 min with 0.1% triton. Cells were incubated with primary antibodies diluted in a blocking buffer overnight at 4°C. After washing with PBS and incubated with Alexa-fluor conjugated secondary antibodies (1:500, various, Thermo Fisher) for 1 hr at room temperature before staining with DAPI (1μg/ml, Merck, D9542) and mounting with Prolong Gold antifade (Invitrogen P36934). For paraffin embedding (Figure 4), organoids were seeded onto 0.4 μm transwell inserts (Greiner Bio-One 662641) embedded in 50% matrigel. Organoids were fixed with 4% PFA for 30 min before the membrane was released from the insert using a scalpel and embedded between layers of HistoGel (Epredia HG-4000-012) before paraffin embedding. Samples were deparaffinised and rehydrated using sequential passes through xylene (x3) then ethanol (100%, 70%, 50%, 0%) then antigen retrieval undertaken by boiling slides in sodium citrate buffer (10mM sodium citrate, 0.05% tween 20 pH 6.0) for 10 minutes. Samples were blocked using 1% BSA, 0.1% tween then incubated with primary antibodies overnight and Alexa-fluor conjugated secondary antibodies (1:500, various, ThermoFisher) for 1 hr at room temperature before staining with DAPI, and mounting with prolong gold antifade. Imaging was undertaken using a Zeiss LSM880 with airyscan and Zen black software.

For whole-mount immunostaining of organoids and 2D AT1-like cells (Figures 1 and 2), the matrigel was completely removed from the cultured organoids using cell recovery solution (Corning, 354253) then fixed with 4% PFA for 30 min on ice. After rinsing in PBS washing solution containing 0.2% (v/v) Triton X-100 and 0.5% (w/v) BSA, the samples were incubated in permeabilization/blocking solution containing 0.2% (v/v) Triton X-100, 1% (w/v) BSA, and 5% normal donkey serum (Stratech Scientific, 017-000-121-JIR) in PBS, overnight at 4°C. Samples were then incubated with primary antibody at 4°C overnight, washed and incubated with secondary antibody for 1 hour at room temperature. Nuclei were counterstained using DAPI. Prior to the organoid imaging, step-wise treatments of 10%, 25%, 50%, and 97% (v/v) 2’−2’-thio-diethanol (TDE, Merck, 166782) were followed for clearing. Images were taken using a Leica SP8 confocal microscope or Zeiss LSM880 with airyscan and Zen black software.

For lysosomal fluorescence, organoids incubated with Lysotracker at 37°C for 2 hr (Cell Signaling Technologies, 8783S) and immediately imaged under a fluorescence microscope.

## Quantitative RT-PCR

Total RNA was isolated using an RNeasy kit (Qiagen, 74004) including an optional DNase digestion step. Typically 500 ng RNA was used as the starting template to create cDNA using a high capacity cDNA reverse transcription kit (Applied Biosystems, 4368814) and heating samples to 25°C for 10 min, 37°C for 2 h and 85°C for 5 min. RT-qPCR was undertaken in 96 well plates using 4.5μl 1:10 cDNA and 10.5 μl of a master mix containing (Sigma-Aldrich, S4438) or SYBR Green PCR Master Mix (Applied Biosystems, 4309155). Primer sequences are listed in Supplementary Table 1. Plates were run on a BioRad RT PCR machine typically using the following programme: 95°C 2 min, 40x (95°C for 30 sec, 55°C for 30 sec, 72°C for 30 sec), 95°C for 30 sec.

## Bulk RNA-sequencing

For AT2 organoid bulk RNA-sequencing, RNA libraries of four biological lines of AT2 organoids at early (P1,4,6,7) and late (P12,13,16,17) passage were generated using an RNeasy kit (Qiagen, 74004) including the optional DNAse digestion step. The quality of the RNA libraries was validated on Agilent 2200 Tapestation before sequencing on an Illumina NovaSeq 6000 at Novogene (novogene.com). A comparison between the RNA sequencing data of AT2 organoids and publicly available data of fetal organoid types and other alveolar cell types was performed: fetal early tip progenitor organoids GSM5393370 and GSM5393371; fetal late tip progenitor organoids^11^ GSM5393372 and GSM5393373; PSC-iAT2s^12^ GSM5578511, GSM5578512 and GSM5578513; cultured adult AT2 cells^12^ GSM5578508, GSM5578509 and GSM5578510; freshly isolated adult AT2 cells^13^ GSM2537127, GSM2537128 and GSM2537129. The raw RNA sequencing data was run by a bioinformatics pipeline, nf-core/rnaseq^14^. A list of differentially expressed genes was extracted using the counted reads and R package edgeR^15^ version 3.40.2. GO biological processes term enrichment, KEGG pathway, and gene set enrichment analysis were performed using DAVID^16^ and R package fgsea^17^ packages. Sequencing data have been deposited at GEO: GSE237359.

## Electron microscopy

Whole organoids were fixed with 2% PFA, 2.5% glutaraldehyde, 0.1 M cacodylate buffer, pH 7.4. Organoids were secondarily fixed with 1% osmium tetroxide/1.5% potassium ferrocyanide and then incubated with 1% tannic acid in 0.1 M cacodylate buffer to enhance membrane contrast. Organoids were washed with water before being dehydrated using increasing percentages of ethanol (70,%, 90%, 100%). Samples were embedded in beam capsules in CY212 Epoxy resin and resin cured overnight at 65°C. Ultrathin sections were cut using a diamond knife mounted to a Reichart ultracut S ultramicrotome. Sections were collected onto piloform-coated slot grids and stained using lead citrate. Sections were viewed on a FEI Tecnai transmission electron microscope at a working voltage of 80 kV.

## Bioinformatic and statistical analysis

Statistical analysis of CRISPR screen sequencing data was performed using MAGeK^18^. Of note, the calculated significance of a gene is not necessarily directly proportional to its biological significance; relative gRNA efficiency and lethal phenotypes generated from knockdowns may preclude enrichment of other genes of functional relevance.

Data expressed as mean ± standard deviation (SD) or standard error of mean (SEM) from at least three independent experiments. Each statistical test is described in the figure legends. Graphpad Prism software (version 9.5.1) was used for statistical analysis and data visualisation.

## Code Availability

No new code was generated for use in this manuscript. Any additional information required to re-run the code and repeat the analyses reported can be requested from the corresponding authors.

